# HDAC6 Regulates Radiosensitivity of Non-Small Cell Lung Cancer by Promoting Degradation of Chk1

**DOI:** 10.1101/2020.02.10.942573

**Authors:** Niko Moses, Mu Zhang, Jheng-Yu Wu, Chen Hu, Shengyan Xiang, Agnes Malysa, Hyejeong Jang, Wei Chen, Wenlong Bai, Gerold Bepler, Xiaohong Mary Zhang

## Abstract

We previously discovered that HDAC6 regulates the DNA damage response via modulating the homeostasis of a DNA mismatch repair protein, MSH2, through HDAC6’s ubiquitin E3 ligase activity. Here, we have reported HDAC6’s second E3 ligase substrate, a critical cell cycle checkpoint protein, Chk1. We have found that HDAC6 and Chk1 directly interact, and that HDAC6 ubiquitinates Chk1 *in vivo*. Typical Chk1 protein levels fluctuate, peaking at G2 and subsequently resolving via the ubiquitin-proteasome pathway. However, in HDAC6 knockdown cells, Chk1 is constitutively active and fails to resolve post-ionizing radiation (IR), leading to increased radiation sensitivity. Upon IR treatment, a greater proportion of HDAC6 knockdown cells accumulated at G2/M phase when compared with control cells. Depletion or inhibition of Chk1 in HDAC6 knockdown cells renders those cells radiosensitive, suggesting that persistently high level of Chk1 could lead cells to arrest at G2/M phase and eventually, apoptosis. Clinically, we found that high levels of phosphorylated Chk1 (p-Ser317) are associated with a better overall survival in a cohort of non-small cell lung cancer (NSCLC) patients, suggesting a link between active Chk1 and lung cancer development. Overall, our results highlight a novel mechanism of Chk1 regulation at the protein level, and a possible strategy for sensitizing NSCLC to radiation via inhibiting the activity of HDAC6’s E3 ligase.

---

Histone Deacetylase 6 is a class IIb HDAC, whose dissimilarity from conventional class I HDACs lies in its primarily cytoplasmic localization. HDAC6’s canonical deacetylation targets include cortactin, acetylated α-tubulin (1), and HSP90. HDAC6 is further set apart from all HDACs in that it contains two catalytic domains, termed DAC1 and DAC2. HDAC6 was first identified as a deacetylase of α-tubulin in a study that described its regulation of cell migration and motility (1), opening up the floodgates to a wave of subsequent studies detailing the oncogenic role that HDAC6 plays in a variety of tumor types, including breast cancer (2), ovarian cancer (3), AML (4), and glioblastoma (5). Our unpublished tissue microarray data also demonstrates that HDAC6 is upregulated across all three subtypes of non-small cell lung cancer (NSCLC), the second most commonly diagnosed cancer across both sexes that is responsible for the greatest number of cancer-related deaths annually. Further clinical innovation is needed to improve outcomes for NSCLC patients.

HDAC inhibition as an anticancer therapeutic strategy has been gaining traction over the last two decades, and has seen major success in the FDA approval of pan-HDAC inhibitors Vorinostat and Romidepsin for treatment of cutaneous T cell lymphoma (CTCL) (6). It has been widely observed that pan-HDAC inhibitors can promote growth arrest, differentiation, and apoptosis in tumor cells with minimal off-target toxicity to the surrounding normal tissue (7). These studies have promoted investigation into the particular HDACs responsible for the differential response between transformed tissue and normal tissue, as inhibition of these HDAC isotypes will enhance tumor responsiveness to intervention, while further mitigating off-target effects of the treatment (8,9). In particular, HDAC6 knockout mice develop normally, which makes HDAC6 an ideal target in cancer treatment (9). HDAC6-specific inhibition has been tested both preclinically and clinically, and a trend has emerged for combining HDAC6 inhibition with DNA damaging agents (10-14). Currently there are two HDAC6-specific inhibitors in Phase I/II clinical trials, ACY-1215 and ACY-241, but much remains to be discovered concerning how HDAC6 interacts with DNA damage response (DDR) proteins and how HDAC6 inhibition impacts tumor DDR activity. In 2012, our group found that knockdown of HDAC6 in NSCLC cell lines sensitized these cells to cisplatin treatment (15). Our current study is the product of linking and expanding the enhanced cisplatin sensitivity of HDAC6 knockdown cells and our 2014 study, which describes E3 ubiquitin ligase activity contained in HDAC6’s DAC1 domain (16). We identified DNA mismatch repair protein MSH2 as a target of HDAC6’s novel E3 ligase activity and deacetylase activity, and found that loss of HDAC6 and subsequent increase of MSh2 rendered cells more sensitive to 6-TG treatment. However, we suspect that HDAC6 may play a more overarching role in the regulation of the DDR via ubiquitination of currently unidentified targets.

Chk1 is a Ser/Thr kinase activated by ATR in response to a variety of DNA aberrations, one of which being the resected ends of a DNA double-strand break (17). The fluctuation of total Chk1 protein levels peaking at S/G2 and decreasing as the cell prepares to re-enter G1 is well documented (18), and multiple E3 ligases have been identified that contribute to Chk1 degradation (19-21). It is fairly common for central DDR proteins to act under the tight control of multiple E3 ligases (22), p53 being the most notable example (23), which suggests that the complete repertoire of Chk1-targeting E3 ubiquitin ligases has yet to be uncovered. In this report, we propose HDAC6 as a candidate Chk1 E3 ubiquitin ligase, as we have confirmed that HDAC6 and Chk1 interact and that HDAC6 can ubiqutinate Chk1 *in vivo*. We have also found that in addition to DNA-damaging chemotherapeutics, genetic ablation of HDAC6 can increase the efficacy of ionizing radiation. Further analysis of irradiated NSCLC cells revealed that in HDAC6-knockdown cells the stable Chk1 is active, as indicated by phosphorylation on S345 and S317. While elevated Chk1 activity has previously been reported to contribute to tumoral genomic stability (24,25), constitutive Chk1 expression and activation resulting from HDAC6 inhibition appears to serve as a detriment to the ability of irradiated NSCLC tumor cells to withstand ionizing radiation.

## Materials and Methods

### Antibodies, Chemicals, and Reagents

Anti-Flag M2 agarose beads (A2220) and anti-HA agarose beads (A2095) were purchased from Sigma. Anti-Chk1 was purchased from Santa Cruz (sc-56288). Anti-PARP-1 (9532), anti-pChk1S345 (2341), anti-pChk1S317 (2344), anti-γ-H2AX (9718), and anti-pCdc25C (9528S) antibodies were purchased from Cell Signaling Technologies. Cycloheximide (C4859), cisplatin (479306), and imidazole (12399) were purchased from Sigma. Ni-NTA resin (635659) was purchased from Clontech.

### Establishment of HDAC6 knockout cell lines

HDAC6 knockout cells were created using CRISPR/Cas9 (Clustered regularly interspaced short palindromic repeates) method. Briefly, the guide RNA targeting HDAC6 exon 5 (5’-GAAAGGACACGCAGCGATCT-3’) was selected and constructed into LentiCRISPRv2 vector (Addgene plasmid 52961). The HDAC6-KO vector can also express the codon-optimized Cas9 protein as well as puromycin resistance gene. The 293T, H1975, and H157 cells infected with the HDAC6-KO virus were selected for stable clones using puromycin at 1 μg/mL. The HDAC6-knockout clones were screened by anti-HDAC6 (H-300) Western blot analysis.

### Cell Culture

NSCLC cell lines A549, H460, and H1299 cells were obtained from the American Type Culture Collection (ATCC). All cell lines were grown in either Dulbecco’s Modified Eagle’s Medium (DMEM) or Roswell Park Memorial Institute 1640 Medium (RPMI), both with 10% fetal bovine serum, penicillin (100U/mL), and streptomycin (100U/mL). All cells were incubated at 37°C with 5% CO_2_. 293T and 293T HDAC6 KO cells were grown in DMEM. H1299, A549, and H460 cells were grown in RPMI. A549 HDAC6 knockdown and control stable cell lines were kindly provided by Dr. Tso-Pang yao. H460 scramble and HDAC6 knockdown stable cells were generated as previously described (15).

### Generation of H1299 inducible HDAC6 knockdown cells

shRNAs for HDAC6 were ordered from Dharmacon (cat. RHS4696), transfected into H1299 cells, and selected with 1μg/mL puromycin for 1 month. Independent clones were then cosen by induction with 0.1μg/mL Doxycycline (D9891 Sigma) for 3 days, and then subjected to Western Blotting for HDAC6. All cells were cultured with tetracycline-free medium.

### Trypan Blue Exclusion

Upon completion of a radiation time course, cells were trypsinized, suspended in PBS, and the density of the solution was quantified via hemacytometer. A 0.4% solution of trypan blue in PBS was prepared, and 0.1mL of this trypan blue solution was added to 0.1mL of cells in PBS. The trypan blue and cell mixture was incubated for 15 minutes at RT, then loaded into a hemacytometer and manually counted under a microscope. The number of blue-stained cells and the number of total cells were recorded, and viability assessed as: % viable cells = [1.00 – (number of blue cells – number of total cells)] x 100.

### Colony Formation Assay

A549 inducible HDAC6 knockdown cells were treated with doxycycline (0.1μg/mL) for 2 weeks, and subsequently seeded in triplicate (300 cells/mL) into 6-well plates. Cells were incubated overnight at 37°C to allow for adherence to the dishes. The cells were then treated with the indicated dose of radiation and incubated for 13 days. Upon completion of the time course, cells were directly fixed and stained with crystal violed (0.05% w/v, 1% formaldehyde, 1% methanol in PBS) for 20 minutes. Colonies on each plate were scanned and counted using the Cell Counter feature of ImageJ software.

### Immunofluorescence staining

A549 control and stable HDAC6 knockdown cells were stained for γ-H2AX and Cyclin A as previously described (26).

### Constructs and transfection

The GST-tagged HDAC6 deletion mutant constructs were synthesized as previously described (16). The Myc-Chk1, Myc-Chk1(1-264) and Myc-Chk1 (265-476) plasmids were generous gifts from Dr. Youwei Zhang (27). Flag-Chk1 was purchased from Addgene (22894). The plasmids were transiently or stably transfected into cells using Lipofectamine 2000 (Invitrogen).

### GST-pull down assay

BL21 cells harboring the GST or various GST recombinant HDAC6 plasmids were grown to log phase and induced with Isopropyl β-D-1-thiogalactopyranoside (IPTG) for 4hrs. After sonication in STE buffer (10 mM Tris-HCL [pH 8.0], 150 mM EDTA, and 5 mM dithiothreitol) containing 1% sarcosyl (w/v, final concentration), solubilized proteins were recovered by centrifugation and incubated with glutathione-agarose beads in the presence of 3% Triton X-100 (final concentration) for 30 min at 4°C and washed several times with ice-cold phosphate buffer saline (PBS). The resulting beads-bound proteins were then incubated with the cell lysates containing the proteins to be pulled down for 4 hrs at 4°C. The glutathione-agarose beads were washed three times with LS buffer prior to loading onto SDS-PAGE.

### Irradiation

Radiation treatment was performed with an X-ray generating Pantak HF 320 instrument (settings at 320kV, 10mA, @ ∼0.9Gy per minute).

### In Vivo ubiquitination assay

Myc-Chk1 (4μg) was cotransfected with 2μg of Flag-HDAC6 and His-Ub as indicated into 293T cells. Thirty-six hours posttransfection, cells were harvested. Cell lysates were lysed in 6M guanidine buffer (50 mM Na2HPO4, 50 mM NaH2PO4, 10 mM Tris.HCl [pH8.0], 10 mM β-mercaptoethanol, 5 mM imidazole, 0.2% Triton X100) overnight with Ni-NTA beads. The beads were washed with 8M urea buffer (50 mM Na2HPO4, 50 mM NaH2PO4, 10 mM Tris.HCl [pH8.0], 10 mM ◸-mercaptoethanol, 5 mM imidazole, 0.2% Triton X100) for 4 times, then eluted with elution buffer (0.5 M imidazole, 0.125 M dithiothreitol (DTT), 1XSDS loading buffer) for 30 minutes at room temperature.

### Generation of Chk1 inducible knockdown cells

TRIPZ inducible lentiviral shRNA against Chk1 was purchased from Dharmacon (RHS4696 glycerol stock). The TRIPZ plasmid was transfected into 293T cells along with lentiviral packaging and envelope 2nd generation plasmids (Addgene packaging 11263, Addgene envelope 17576). 48 hours post-transfection, the DMEM media was collected from the 293T cells, and 200μL of this media was added to A549 HDAC6 stable knockdown cells at ∼60% confluence in a 6cm dish. These A549 cells were expanded and split into a 10cm dish, at which point doxycycline (0.1 μg/ml) was added. Cells were treated with doxycycline for 2 weeks to induce the expression of the TRIPZ vector, which will co-express shRNA against Chk1 and fluorescent marker TurboRFP. RFP-positive cells were sorted using a Sony SY2300 (Sterile/4-Way) cell sorter incorporated into a Baker SterilGARD II Biological Safety Cabinet, allowing for sterile BSL Class II sorting. This sorting produced the Chk1Tripz pool. Further experiments were conducted after establishment of stable clones with high expression of RFP.

### RT-PCR

Reverse transcriptase-polymerase chain reaction (RT-PCR) assays were performed to measure the expression of mRNA. Cells were washed at least twice with PBS and immediately lysed in Trizol® (AMBION, Catalog number: 15596026). For Mouse tissue samples: Add 1 mL of TRIzol™ Reagent per 50–100 mg of tissue to the sample and homogenize using a homogenizer. Total RNA was then isolated to follow the Trizol method (Invitrogen). Subsequently, 1 ug of RNA was reverse-transcribed using the WarmStart® RTx reverse transcriptase (New England BioLabs, M0380L) and random primer mix (New England BioLabs, S1330S) according to the typical cDNA synthesis protocol. PCR reactions were performed with Taq 2X Master Mix (New England BioLabs, M0270L). The thermocycler conditions for Mouse Chk1 and GAPDH PCR product were as follows: 95°C 30 sec for 1 cycle; 95°C 30 sec, 55°C 60 sec, and 68°C 1 min for 40 cycles; final extension 68°C 5 min for 1 cycle. The same thermocycler conditions were used for Human Chk1 and GAPDH. The following PCR primers were used for RT-PCR: Human Chk1-forward: 5’-ATGCTCGCTGGA GAATTGC-3’; Human Chk1-reverse: 5′-ATA AGGAAAGACCTGTGCGG-3’; Human GAPDH-forward: 5′-GGAGCGAGATCCCTC CAAAAT-3′; Human GAPDH-reverse: 5′-GGC TGTTGTCATACTTCTCATGG-3.′ Mouse Chk1-forward: 5’-CTTTGGGAGAAG GTG CCTAT-3’; Mouse Chk1-reverse: 5′-ATG CCG AAATACCGTTGC-3’; Mouse GAPDH-forward: 5′-GTTGTCTCCTGCGACTTCA-3′; Mouse GAPDH-reverse: 5′-GGTGGTCCA GGGTTTCTTA-3.′ The PCR products were then loaded onto agarose gel with ethidium bromide. After gel electrophoresis, the PCR products were visualized under the UV light.

### TMA-1 Composition

TMA-1 consists of 252 early stage NSCLC patients and a series of normal tissue controls: larynx, stomach, trachea, esophagus, liver, breast, lymph node and normal adjacent lung. Triplicate tissue cores were put onto the three tissue array blocks (TMA 1-7, 1-8 and 1-9) to account for differences that could arise in tissue staining.

### Patient Cohort

The single institution cohort of 187 early-stage NSCLC patients treated by surgery alone has been described (28). Cohort demographic and clinical characteristics are summarized in Table 1. The median follow-up time for overall survival (OS) was 69.0 months. The study was approved by the IRBs of the Wayne State Unviersity and conformed to the Helsinki declaration.

**Table 1.** Demographic and clinical characteristics of the cohort

| Variable |  | NSCLC<br>(n=187) |
| --- | --- | --- |
| Age | Median | 70.4 |
|  | Range(min, max) | (45.9,84.4) |
| Sex | F | 86 (46%) |
|  | M | 101 (54%) |
| Race | African American | 6 (3%) |
|  | Caucasian | 179 (96%) |
|  | Other | 2 (0.01%) |
| PS | 0 | 128 (68%) |
|  | 1 | 48 (26%) |
|  | Unknown | 11 (6%) |
| Stage | IA | 85 (45%) |
|  | IB | 102 (55%) |
| Histology | Adeno | 78 (42%) |
|  | BAC | 18 (10%) |
|  | Large Cell | 20 (11%) |
|  | LNE | 3 (2%) |
|  | Squamous | 68 (36%) |
| Smoker | Never | 11 (6%) |
|  | Active | 49 (26%) |
|  | Quit | 113 (60%) |
|  | Unknown | 14 (7%) |

### Tumor Samples

Tissue microarrays (TMA) consisting of formalin fixed paraffin embedded (FFPE) NSCLC patient tumor tissue were analyzed and each patient had three distinct cores. TMAs were stained using a standard immunofluorescence protocol and primary antibody concentrations were optimized individually at two institutions (Moffitt Cancer Center, Tampa, FL and Karmanos Cancer Institute, Detroit, MI). Slides were acquired and scored using Automated Quantitative Analysis (AQUA) (PM2000 version 2.3.4.1, Genoptix, Carlsbad, CA). Signal intensities for nuclear, cytoplasm, and/or membrane compartments were scored.

### AQUA Analysis

To perform phosphorylated S317 Chk1 AQUA staining, TMA-1 replicates (TMA 1-7, 1-8 and 1-9) were subjected to a series of xylenes, graded alcohol and distilled water washes. Antigen retrieval was done using 1X PT Module buffer solution (Thermo Scientific, Catalog#: TA-250-PM4x) and Decloaking Chamber Pro (Biocare Medical, Catalog#:DC2002), with the following protocol: 125oC for 30 seconds, 90oC for 10 seconds and cool to room temperature. Tissue arrays were blocked with 5% Normal Goat Serum (NGS) (Invitrogen, Catalog#: PCN5000)+ 0.1% Tween 20 (T20)+ 1x PBS (PBS) for 1 hour at room temperature. Tissue arrays were then stained with 1:25 Chk1 S317 anti-rabbit (Novus Biologicals, Catalog#: NB100-92499) and pan-cytokeratin AE1/AE3 anti-mouse (1:200, DAKO Cytomation, Catalog#: M3515) using the following diluent: 1% NGS+ 0.1% T20+PBS.

Tissue arrays were washed with the following wash buffer: 1%NGS+0.4%TritonX+PBS and incubated with secondary antibodies for 1 hour at room temperature with anti-mouse IgG (H+L) Alexa Fluor 555 (1:200, Thermo Fisher, Catalog# A-21422) or anti-rabbit IgG (H+L) Alexa Fluor 555 (1:200, Thermo Fisher, Catalog#: A-21428) diluted in wash buffer. Slides were mounted using Prolong Gold anti-fade reagent with DAPI (Thermo Fisher, Catalog#: P36931). Measuring levels of phosphorylated S317 Chk1 in nuclear and cytoplasmic compartments was performed using AQUA technology (PM-2000, version 2.3.4.1, Genoptix, Carlsbad, CA).

### Statistical Analysis

AQUA measurements were standardized into mean 0 and standard deviation 1. Replicate cores were then averaged for each patient. The primary clinical endpoint for analysis was overall survival (OS), defined as the time interval from the date of resection to death. A search for optimal cut-off of AQUA measurements was performed using Cox model adjusted for tumor stage and age. An optimal cut-off was defined as the one with lowest Cox adjusted p value constrained that the minimum sample size from the resulted high/low groups was larger than 50. KM plot was provided for the high and low maker groups. All p-values were two-sided with a significance level of 0.05. The 95% CI for median survivals in the subgroups were calculated with log-log method. All calculations were performed with R statistical program (R Project for Statistical Computing, Version 3.4.3, Vienna, Austria).

## Results

### HDAC6 influences Chk1 protein stability

HDAC6 is unique amongst the HDAC family members in that it possesses both deacetylase activity and E3 ubiquitin ligase activity. While a myriad of substrates have been identified for HDAC6’s deacetylase activity, to date MSH2 is the only reported target of HDAC6’s E3 ubiquitin ligase activity (16). We suspected that there may be additional targets of this activity, specifically proteins involved in the DDR.

Our group has previously generated A549, H157, and H1975 HDAC6 knockout cells using the CRISPR/Cas9 system. We began by probing these cell lines for HDAC6, acetylated α-tubulin (a downstream target of HDAC6 deacetylase activity), Chk1, and GAPDH. Across all three cell lines, successful HDAC6 knockdown corresponded with an increase in baseline Chk1 protein [Figure 1A, lanes 1-6]. Inducible HDAC6 knockdown cell lines H1299 and A549 were pre-treated with doxycycline for two weeks, and these cells exhibited complete (H1299) and partial (A549) HDAC6 knockdown accompanied by an increase in Chk1 protein levels in both cell lines [Figure 1A, lanes 7-10]. Mouse embryonic fibroblasts (MEFs) obtained from C57Bl/6 wild-type and HDAC6 knockout mice were also tested, and we observed the same increase in Chk1 protein in the HDAC6 knockout MEFs compared to the controls [Figure 1A, lanes 11-12]. Finally, we harvested a variety of organ samples from our C57Bl/6 wild-type and HDAC6 knockout mice, and across every tissue type we tested an absence of HDAC6 protein corresponded with a higher basal level of Chk1 than observed in HDAC6-competent mice [Figure 1A, lanes 13-24].

**Figure 1:**
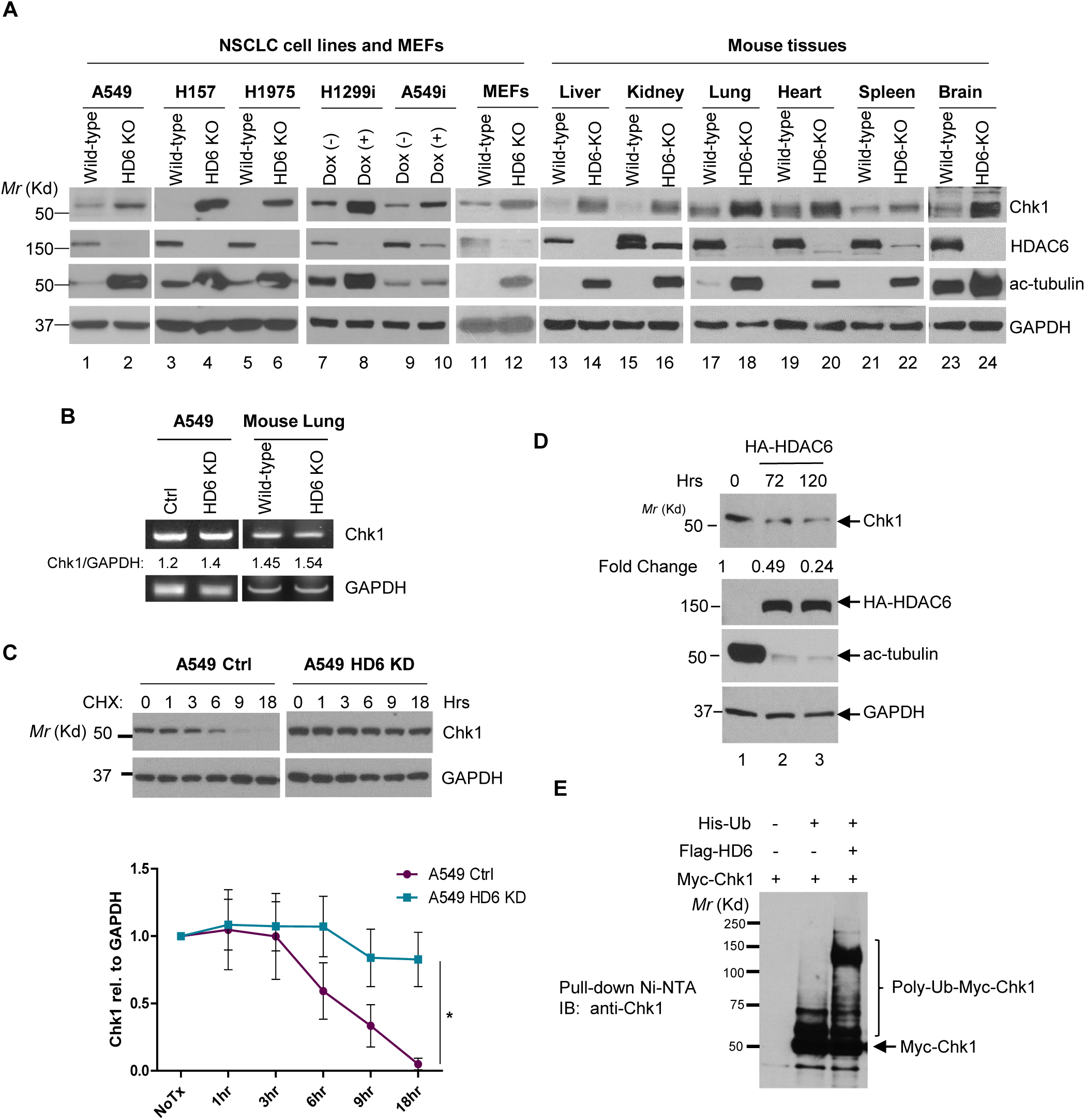
HDAC6 regulates Chk1 protein stability. **A.)** (From left to right) A549 HDAC6 KO cells generated with the CRISPR-Cas9 system. H157 and H1975 HDAC6 KO cells generated with the CRISPR-Cas9 system. H1299 and A549 inducible HDAC6 knockdown cells (termed H1299i and A549i, respectively) pre-treated with doxycycline for two weeks. Mouse embryonic fibroblasts (MEFs) harvested from age-matched wild-type and transgenic HDAC6 KO mice (both from a C57Bl/6 background). Liver, kidney, lung, heart, spleen, and brain tissue harvested from age-matched wild type and transgenic HDAC6 KO mice (both from a C57Bl/6 background). All cell lines and tissues were lysed and analyzed via Western Blot for Chk1, HDAC6, acetylated tubulin, and GAPDH expression. **B.)** RT-PCR was used to determine whether HDAC6 knockdown influences Chk1 mRNA levels in A549 control and HDAC6 stable knockdown cells, as well as WT and HDAC6 knockout murine lung tissue. **C.)** (Above) A549 stable knockdown cells were treated with 10μg/mL cycloheximide (CHX), harvested at the indicated timepoints, and analyzed via Western blot. Representative Western blot of Chk1 and GAPDH from the trials used to determine Chk1 half-life. (Below) The average intensity of Chk1 relative to GAPDH expression from three independent experiments was obtained (via ImageJ) and graphed. **D.)** 293T HDAC6 knockout cells were plated, and 24 hours later were transfected with 2.4μg HA-tagged HDAC6. Control cells were treated with transfection reagent PEI for 24 hours. HA-HDAC6-transfected cells were harvested at the indicated timepoints and probed for the indicated proteins. Fold-change in Chk1 expression was evaluated via ImageJ. **E.)** Mammalian expression vectors containing Myc-Chk1, Flag-HDAC6, and His-Ub were transfected into HEK-293T cells. Cells were incubated for 48 hours, harvested, and passed through a Ni-NTA column to pull down for His-Ub. Bound proteins were subsequently eluted from the columns, run on an SDS-PAGE gel, and probed for Chk1.

To determine whether Chk1 expression is being regulated at the protein level or the mRNA level, RT-PCR for Chk1 was performed in our A549 stable knockdown cells as well as wild-type and HDAC6 KO murine lung tissue. Knockdown or knockout of HDAC6 did not influence Chk1 mRNA content [Figure 1B], so we proceeded by examining the regulation of Chk1 expression at the protein level.

To examine whether the increase of Chk1 in HDAC6-knockdown cells is due to increased protein stability, we treated A549 stable knockdown cells with 10 μg/mL cycloheximide, a protein synthesis inhibitor, over an 18 hour timecourse. Previous studies have reported a Chk1 half-life of ∼4.6 hours in parental cancer cells (20), so this timecourse should model typical Chk1 resolution in the control cells. Indeed, Chk1 degraded as expected in the control cells and was virtually undetectable 18 hours post-cycloheximide treatment. In contrast, Chk1 was exquisitely stable in the HDAC6 knockdown cells [Figure 1C]. This protocol was repeated three times, and Chk1 resolution was exponentially modeled in each cell type [data not shown]. Chk1 half-life in the control cells is ∼5.35 hours, while Chk1 half-life in HDAC6 knockdowns is ∼39.95 hours, indicative of Chk1’s significant stability in the absence of HDAC6. Beginning to explore whether HDAC6 could decrease the level of Chk1, we overexpressed HA-tagged HDAC6 in 293T HDAC6 knockout cells. Our untransfected cells maintained high acetyl-tubulin levels, and we used their expression of Chk1 as a control. When HDAC6 was overexpressed for 72 and 120 hours, Chk1 protein levels were reduced by half and three-quarters, respectively as compared to the control [Figure 1D], suggesting that HDAC6 is able to down-regulate the level of Chk1.

Our group previously uncovered ubiquitin E3 ligase activity within the DAC1 domain of HDAC6, which regulates MSH2 stability via ubiquitination both in vitro and in vivo (16). The observed association between loss of HDAC6 and increased Chk1 stability may indicate a role for HDAC6 in the ubiquitination and subsequent degradation of Chk1. To test this possibility, we performed the Ub assays under the denatured conditions. As shown in Figure 1E, overexpression of HDAC6 in 293T cells greatly increased the level of ubiquitinated Chk1, suggesting that HDAC6 is able to ubiquintinate Chk1 in vivo.

### HDAC6 and Chk1 physically interact

As discussed above, MSH2 is currently the only published target of HDAC6’s E3 ligase, whose protein levels are regulated by HDAC6 via sequential deacetylation and ubiquitination (16). HDAC6 directly interacts with MSH2, so it stands to reason that Chk1 may directly interact with HDAC6 as well. Specifically, we hypothesize that HDAC6 directly interacts with and ubiquitinates Chk1, leading to degradation of Chk1. If this hypothesis is correct, enhanced Chk1 stability in HDAC6 knockdown and knockout cells would be due to the elimination of a critical E3 ligase for Chk1 degradation.

We began by assessing whether HDAC6 and Chk1 interact. As shown in Figure 2A and 2B, reciprocal co-immunoprecipitation assays indicate that Flag-tagged Chk1 and HA-tagged HDAC6 interact in 293T cells. To investigate whether this HDAC6 and Chk1 interaction is under a normal condition, we performed coimmunoprecipitation for endogenous proteins in 293T cells, and found that endogenous Chk1 was able to interact with HDAC6 [Figure 2C]. To ensure that this is a direct interaction and not the result of cofactor participation, bacterially purified protein was utilized. As shown in Figure 2D, glutathione-agarose bound GST-HDAC6 was able to pull-down His-Chk1, suggesting that HDAC6 and Chk1 physically interact with each other.

**Figure 2:**
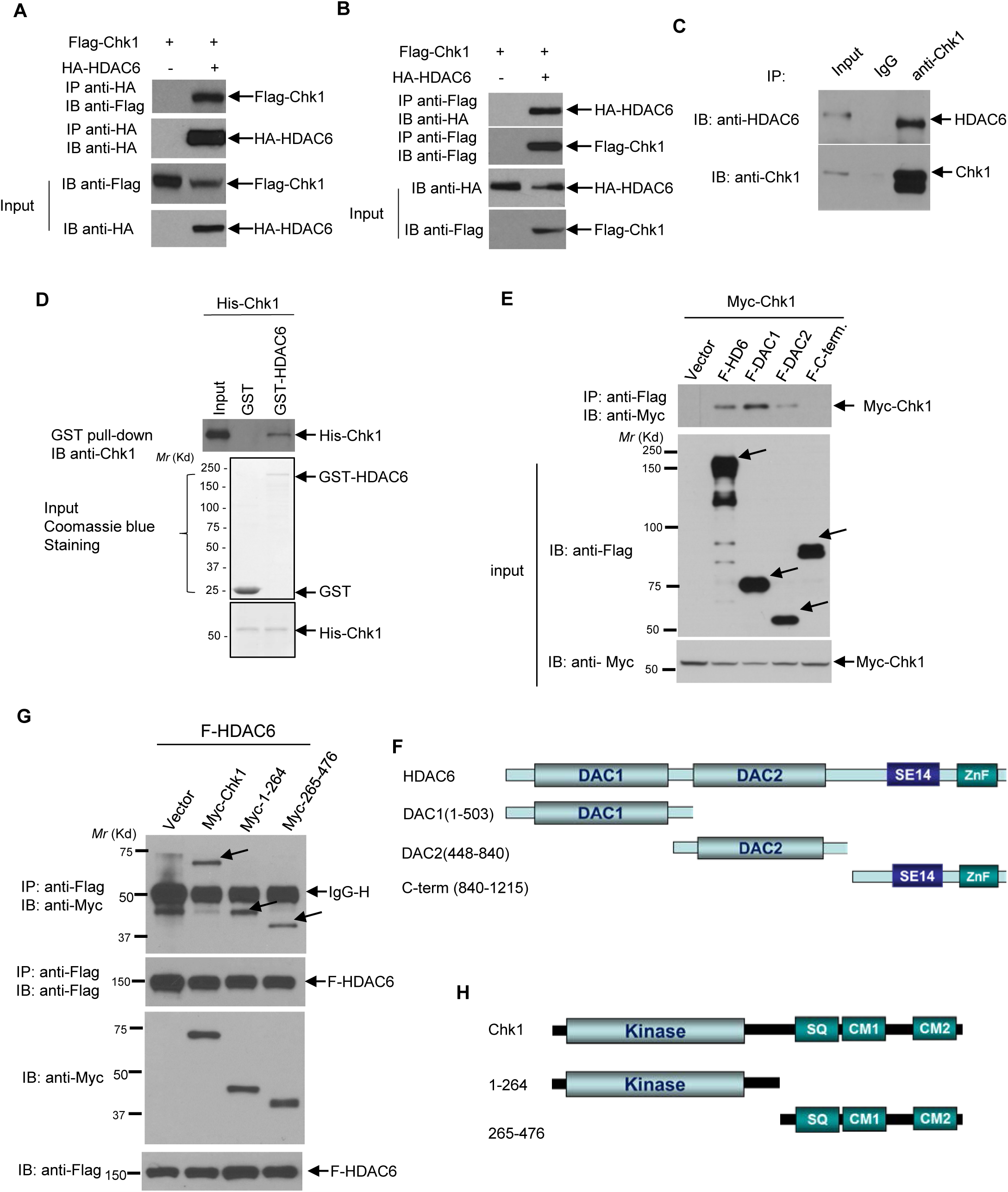
HDAC6 and Chk1 physically interact. **A**,**B.)** Mammalian expression vectors containing Flag-Chk1 and HA-HDAC6 were transfected into HEK-293T cells with PEI. 48 hours after overexpression, cells were harvested in lysis buffer, incubated with either HA-coated (A) or Flag-coated (B) agarose beads, and the resultant immunoprecipitated protein was run on an SDS-page gel and probed for the reciprocal tag. **C.)** 293T lysates were probed with anti-Chk1 antibody complexed with protein A/G beads, the beads were washed, and the resulting milieu probed for HDAC6 to detect an endogenous interaction between Chk1 and HDAC6. **D.)** His-Chk1 was overexpressed in *E. coli*. His-Chk1 was purified with Ni-NTA agarose beads. Then, GST and GST-HDAC6 were overexpressed in *E. coli*, and GST-tagged protein was pulled-down and purified by glutathione-agarose. Purified His-Chk1 was incubated with either glutathione agarose-bound GST or GST-HDAC6, and then bound proteins were eluted. The samples were subjected to SDS-PAGE and Western blot analysis. **E.)** The indicated Flag-tagged HDAC6 deletion mutant constructs were transfected into 293T cells along with Myc-Chk1. 48 hours later, cells were lysed, and lysates were pulled down for Flag. **F.)** Schematic of the Flag-tagged HDAC6 deletion mutant constructs used for the coimmunoprecipitation in (C). **G.)** The indicated Myc-tagged Chk1 deletion mutant constructs were transfected into 293T cells along with Flag-HDAC6. 48 hours later, cells were lysed, and lysates pulled down for Flag. **H.)** Schematic of the Myc-tagged Chk1 deletion mutant constructs used for the coimmunoprecipitation in (E).

Assured that HDAC6 and Chk1 can interact in vitro and in an in vivo, we sought to determine which region of HDAC6 is responsible for the interaction with Chk1. From N-terminus to C-terminus, HDAC6 consists of an E3 ligase-containing DAC1 catalytic domain, the deacetylase-containing DAC2 catalytic domain, a cytoplasmic-anchoring SE14 motif, and a ubiquitin-interacting zinc finger motif (ZnF). Flag-tagged deletion mutants were constructed as indicated in the Figure 2F scheme. These mutant proteins were overexpressed along with Myc-Chk1 in 293T cells. As shown in Figure 2E, the DAC1 domain (1-503) displayed the strongest interaction with Myc-Chk1, followed by full length HDAC6 (1-1215). There was minimal DAC2 domain binding (448-840), while the C-terminal mutant (840-1215) failed to interact with Myc-Chk1. Both of the deletion mutant constructs that displayed a strong interaction with Myc-Chk1 shared a common feature; they contain the DAC1 E3 ligase domain [Figure 2F].

In the reciprocal experiment, we found that the full-length Chk1 along with C-terminal and N-terminal deletion mutants were able to interact with Flag-HDAC6 [Figure 6G and 6H]. These results may imply a plausible multi-faceted interaction between Chk1 and HDAC6 (29), but the exact regions that participate in this interaction remain unidentified.

**Figure 3:**
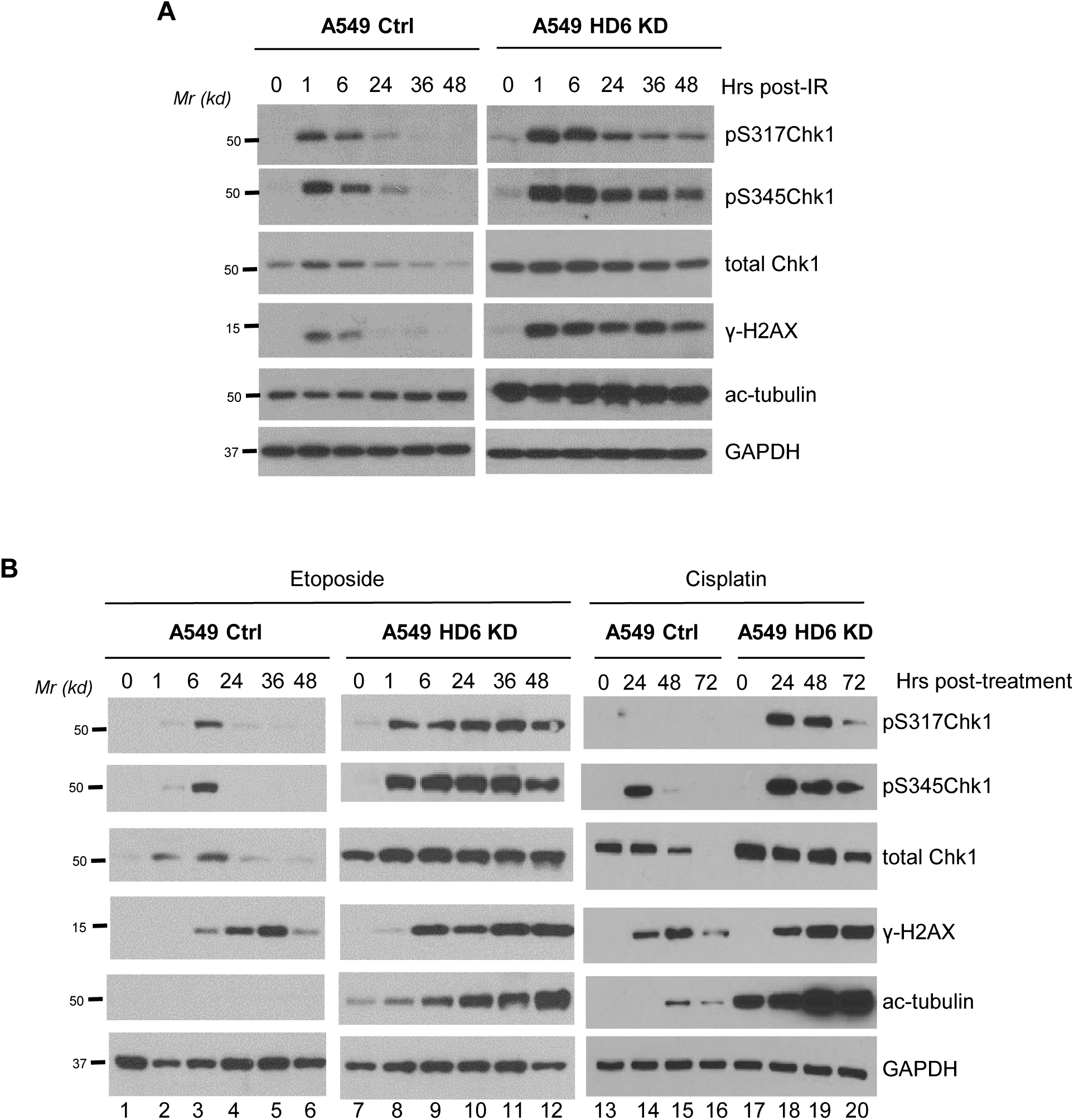
Chk1 is constitutively active in HDAC6 knockdown cells post-DNA damage. **A.)**A549 control and HDAC6 knockdown cells were irradiated with a dose of 10Gy, harvested at the indicated timepoints, and lysates were analyzed via Western blot. **B.)** A549 control and HDAC6 knockdown cells were treated with 20μM Etoposide. Cells were harvested at the indicated timepoints, and lysates were analyzed via Western blot. (left two panels, lanes 1-12) A549 control and HDAC6 knockdown cells were treated with 10 μM Cisplatin. Cells were harvested at the indicated timepoints, and lysates were analyzed via Western blot (right panel, lanes 13-20).

**Figure 4:**
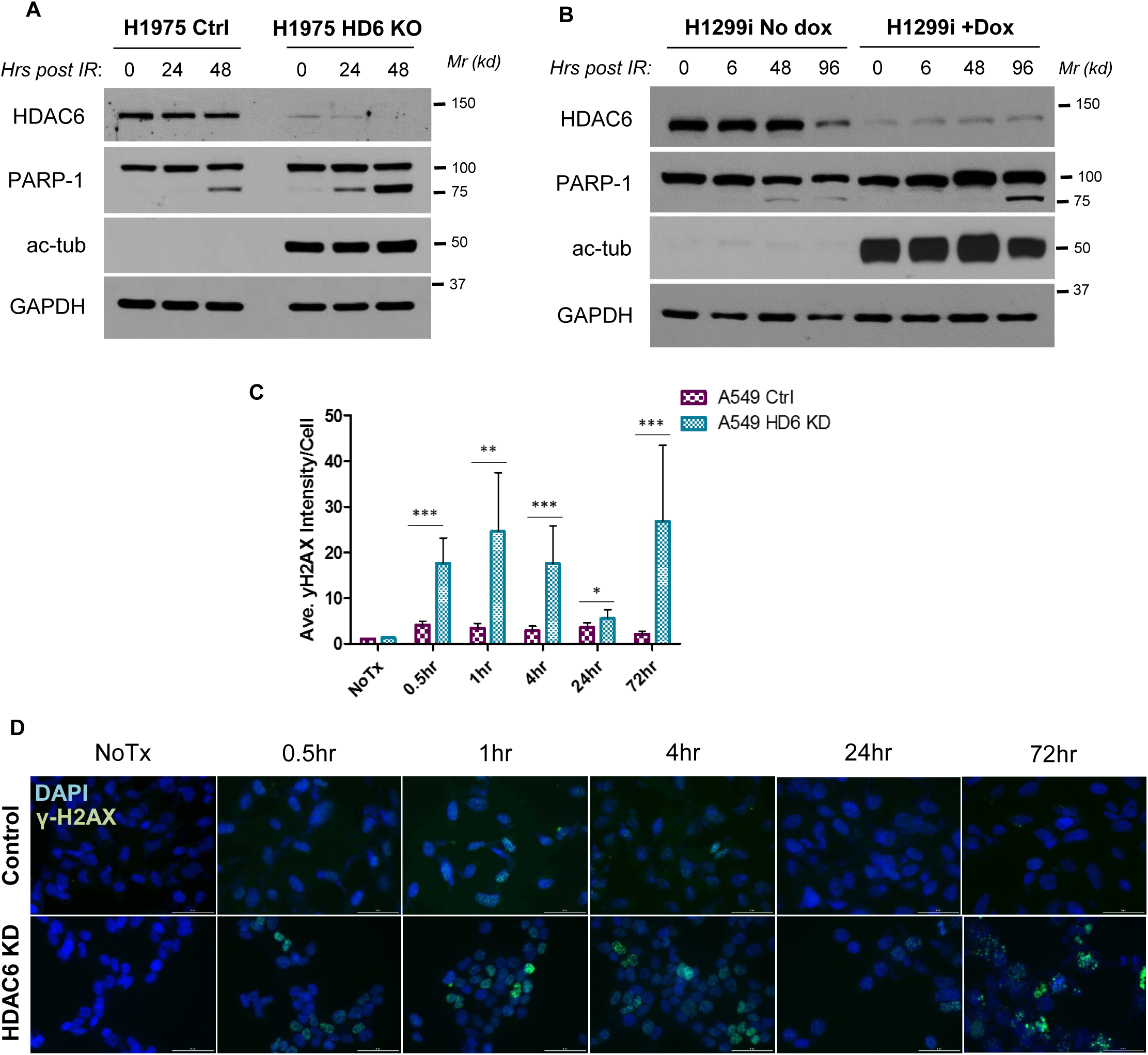
HDAC6 knockdown cells accumulate damage post-radiation. **A.)** H1975 HDAC6 knockout cells were irradiated with 10Gy. Cells were harvested at the indicated timepoints, and analyzed via Western blot for the indicated damage markers and controls. **B.)** H1299i cells were treated and analyzed as described in (A). **C.)** A549 stable knockdown cells were treated with 5Gy radiation. Immunoflourescence staining was performed for γ-H2AX, and mounted on a drop of DAPI-containing mounting media. Intensity of foci was quantified using ImageJ. Student’s t test, ***p<0.0001, **p<0.001, *p<0.05. **D.)** Representative images of the data graphed in (C).

**Figure 5:**
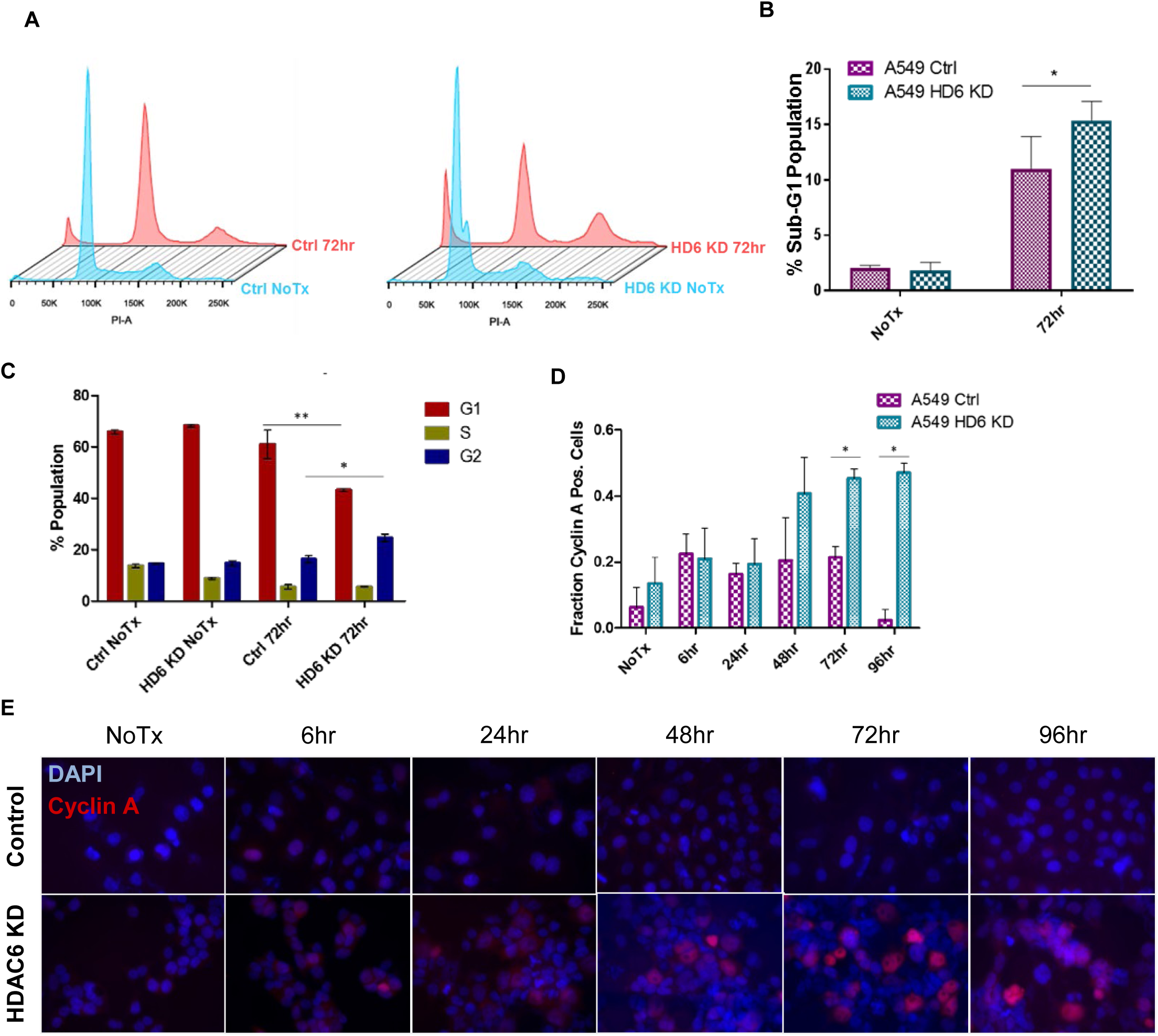
HDAC6 knockdown cells arrested at G2 phase. **A.)** A549 stable HDAC6 knockdown cells were irradiated with 10Gy, incubated for 72 hours, harvested and ethanol fixed, and stained with PI. Cells were analyzed via flow cytometry. **B.)** Analysis of cell cycle distribution from the experiment described in (F). **C.)** Analysis of the fraction of sub-G1 cells present in the experiment described in (F). **D)** A549 stable knockdown cells were treated as described in (D), and immunofluorescence for Cyclin A was conducted. Results of Cyclin A positivity from three biological replicates, with significance assessed using student’s t test, with *p<0.005. **E.)** Representative images of the data graphed in (D).

**Figure 6:**
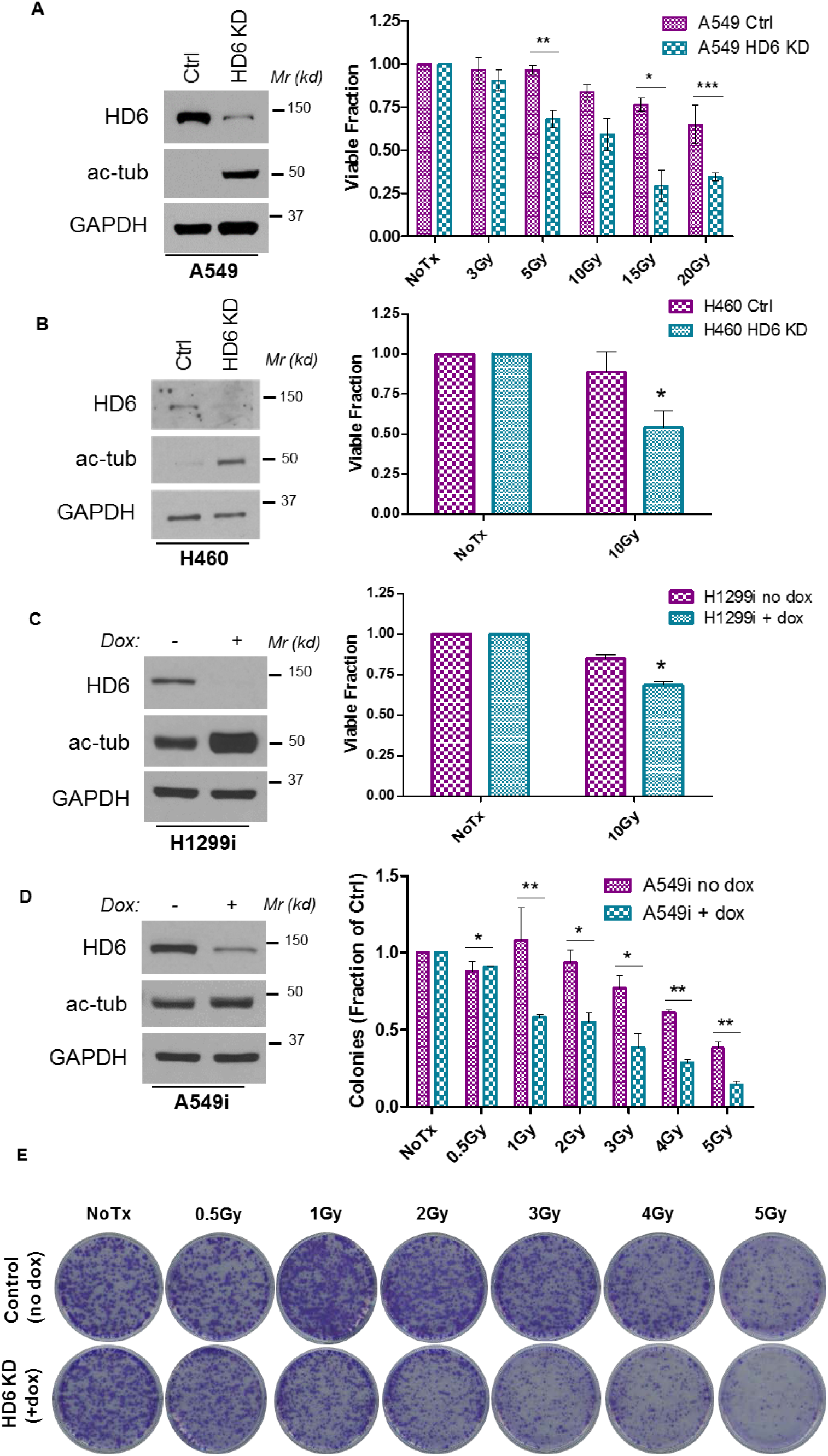
HDAC6 knockdown sensitizes NSCLC cells to radiation. **A.)** (Left) Western blot confirming HDAC6 knockdown in A549 cells. (Right) 120 hours post-radiation, A549 stable knockdown cells were suspended in trypan blue. The number of unstained cells (viable), stained cells (non-viable), and total numbers were recorded. Three biological replicates are graphed. Single sample t-test, *p=0.0122, **p=0.0099, ***p=0.0021. **B.)** (Left) Western blot confirming HDAC6 knockdown in H460 cells. (Right) H460 stable HDAC6 knockdown cells were either left untreated, or treated with 10Gy IR. 120 hours later, trypan blue staining was conducted as described in (A). Single sample t test, mean of 87.77037, *p=0.0154. **C.)** (Left) Western blot confirming inducible HDAC6 knockdown in H1299i cells pre-treated with doxycycline for two weeks. (Right) H1299i cells were either left untreated, or treated with 10Gy IR. 120 hours later, trypan blue staining was conducted as described in (A). Single sample t test, mean of 92.00285, *p=0.0002. **D.)** A549i cells were plated in 6 well plates at a concentration of 350 cells/well, incubated for 24 hours, and irradiated with the indicated dose. 14 days later, cells were stained with crystal violet. Single sample t test, *p<0.02, **p<0.005. Error bars, S.D. **E.)** Representative images from the experiments performed in (D).

### Chk1 is constitutively active in HDAC6 knockdown cells post-DNA damage

Chk1 basal levels were markedly increased in a panel of HDAC6 knockout cells compared to their control counterparts, so we wanted to explore the damage response phenotype resulting from elevated Chk1 in response to ionizing radiation. We treated A549 stable HDAC6 knockdown cells with 10 Gy of IR and assessed total Chk1, active Chk1, and γ-H2AX as a surrogate for double strand breaks. We were able to confirm Chk1’s enhanced stability in the knockdowns, and found that both pChk1 S317 and pChk1 S345 were readily detectable throughout the timecourse (1, 6, 24, 36, 48 hr). This persistence of active Chk1 occurred in parallel to the inability of these cells to resolve DNA DSBs, as indicated by γ-H2AX persistence [Figure 3A]. This timecourse was repeated in our H1299 inducible HDAC6 knockout cells and we observed the same phenotype; stable and active Chk1, and more persistent γ-H2AX than what we observed in the controls [data not shown].

Given that our current study is partially built upon an enhanced cisplatin responsiveness in HDAC6 knockdown cell lines (15), and have confirmed enhanced radiation sensitivity in these HDAC6 knockdowns, we wondered whether this enhanced Chk1 activation phenotype also occurs upon cisplatin treatment. Extending this line of reasoning begs the question of whether this response is radiation-specific or can be observed upon treatment with a broader spectrum of DNA-damaging agents. We tested the radiomimetic drug etoposide (ETO) in our A549 stable knockdown cells using a 20 μM dose, and found the same Chk1 activation and γ-H2AX persistence phenotype that occurred in our radiation timecourse study [Figure 3B, lanes 1-12]. To test an agent whose primary mechanism of action is not double strand break formation, we conducted a cisplatin treatment timecourse in the A549 stable knockdown cells, and detected the Chk1 activation and γ-H2AX persistence phenotype [Figure 3B, lanes 12-20]. The consistency of this response between cell lines and DNA-damaging agents highlights a potential mechanism of cell death in the absence of HDAC6, specifically in the absence of HDAC6’s E3 ubiquitin ligase activity.

### HDAC6 knockdown cells accumulate damage, increase apoptosis and arrest at G2 phase post-radiation

In HDAC6-deficient cell lines, it appears that both total and active Chk1 are present at higher levels post-DNA damage than in the control cells. Next, we wanted to examine whether Chk1’s activity is associated with sensitivity or resistance to ionizing radiation treatment. We assessed PARP-1 cleavage in our cell lines, a marker of apoptosis. In H1975 stable HDAC6 knockout cells, induction of PARP-1 cleavage was observed at 48 and 72 hours post-radiation. In contrast, PARP-1 cleavage was faintly detectable at 72 hours post-IR control cells [Figure 4A]. Preferential PARP-1 cleavage was also observed in our H1299 inducible HDAC6 knockdown model [Figure 4B], suggesting that depletion of HDAC6 sensitizes NSCLC cells to ionizing radiation.

Double-strand break (DSB) formation was also assessed via γ-H2AX. As histone variant, H2AX within the vicinity of a DSB is rapidly phosphorylated by ATM (30). To visualize the damage in situ, we performed immunofluorescence for γ-H2AX. In both A549 control and HDAC6 knockdown cells, we observed a peak 1 hour post radiation, although the HDAC6 knockdown cells exhibited a significantly greater damage burden compared to the controls [Figure 4C-D]. However, at 24 hours the average intensity of foci per cell had decreased in both groups compared to the intensity of foci at 4 hours [Figure 4C], so it stands to reason that both groups may be able to completely resolve the damage if given adequate time. The timecourse was extended out to 72 hours, where we saw a significant accumulation of foci in the HDAC6 knockdowns compared to the controls [Figure 4C-D]. This pattern of γ-H2AX intensity, decreasing and then increasing again, may be indicative of cells attempting to resolve DSBs and failing (31), or the result of apoptotic cleavage of genomic DNA (32).

Interestingly, at 72 hours the HDAC6 knockdown cells displayed an aberrant nuclear morphology riddled with strong, nonhomogenous foci staining, as well as foci-positive micronuclei [Figure 4D]. These structures can be considered markers of mitotic catastrophe (MC), a method of cell death resulting from premature induction of mitosis (33) prior to the cell completing S or G2 phase. MC is estimated to account for ∼60% of cell death in patients treated with ionizing radiation (34), but can also result from treatment with a DNA-damaging agent (33). Additional hallmarks of MC include an increase in the 4N population due to chromosomal segregation failure, an increase in the sub-G1 population, giant cells, multinucleated cells, and micronuclei (35). We further assessed this phenomenon by analyzing the cell cycle distribution of our A549 stable HDAC6 knockdown cells via Propidium Iodide (PI) staining. We observed an increase in G2 arrest in the HDAC6 knockdown cells 72 hours post-10 Gy compared to the controls and a significantly higher sub-G1 population, the latter population being indicative of cell death [Figure 5A-B]. We confirmed this preferential arrest with immunofluorescence staining for cyclin A, a critical cyclin for late-S and G2-phase cells (36), and revealed a preferential accumulation in HDAC6 knockdown cells at later timepoints of post-irradiation [Figure 5C-E]. The accumulation of G2-phase cells staining positive for Cyclin A led us to ponder whether Chk1, the gatekeeper of S and G2 phases of the cell cycle, is functionally involved in the differential response of control and HDAC6 knockdown cells to radiation.

### HDAC6 depletion mediates the reduction of NSCLC cell viability and growth

We previously found that HDAC6 knockdown preferentially sensitizes cells to cisplatin treatment. This sensitization was presumed to be mechanism-specific, as parallel treatment with paclitaxel did not further sensitize HDAC6 knockdown cells (15). While the interstrand DNA crosslinks induced by cisplatin differs from the single- and double-strand DNA breaks ionizing radiation generates, we suspect that the efficacy of treatment in HDAC6 knockdown cells relies on direct DNA damage.

We assessed the viability of A549 and H460 stable HDAC6 knockdown cells and H1299 HDAC6 inducible knockdown cells via trypan blue exclusion. Our A549 stable HDAC6 knockdown cells were treated with the indicated doses of radiation and incubated for 120 hours, at which point they were harvested and stained with trypan blue exclusion dye [Figure 6A]. We found a significant and dose-dependent reduction in viability in the HDAC6 knockdown cells, and confirmed this reduction in H460 stable HDAC6 knockdown cells and H1299 inducible HDAC6 knockdown cells 120 hours post-10Gy radiation [Figure 6B,C]. These data indicate that regardless of whether HDAC6 knockdown is acute or chronic, its loss contributes to exacerbated reductions in viability in NSCLC models.

We next sought to determine the proliferative capacity of our cells post-radiation via colony formation assay. Initial assessment of the A549 and H460 stable knockdown cells revealed a differential plating efficiency between the controls and the HDA6 knockdowns, making a case for HDAC6 inhibition as a single treatment modality but inconclusive when assessing radiosensitization [data not shown]. To subvert this shortcoming of stable knockdown cell lines, we tested the proliferative capacity of our A549 inducible HDAC6 knockdown cells. These cells successfully corrected for the plating efficiency error, and demonstrated a preferential reduction of proliferative capacity in the HDAC6-knockdown cells at 1, 2, 3, 4, and 5 Gy [Figure 6D-E], suggesting that depletion of HDAC6 causes a reduction of a short-term as well as a long-term survival post-IR.

### Radiosensitivity of HDAC6 knockdown cells is dependent on Chk1 activity

We have observed that HDAC6 knockdown can both promote accumulation of active Chk1 [Figure 3] and sensitize cell lines to radiation-induced damage [Figure 6]. We hypothesize that these two phenomena are linked via a causal relationship, and have established this relationship using both genetic and pharmacological inhibition of Chk1. To genetically inhibit Chk1, we produced an A549 HDAC6 stable and Chk1 inducible knockdown cell line by stably transfecting A549 HDAC6 knockdown cells with a TRIPZ inducible shRNA plasmid targeted against Chk1, referred to hereafter as HDAC6 KD + Tripz. After confirmation of the Chk1 knockdown [Figure 7A], we assessed the proliferative capacity of Chk1Tripz cells compared to A549 HDAC6 stable knockdown cell lines via colony formation assay. Using an escalating dose of IR (1, 2, 3, 4 and 5 Gy), we found that genetic ablation of Chk1 had a protective effect on the ability of these cells to grow post-radiation [Figure 7B-C].

**Figure 7:**
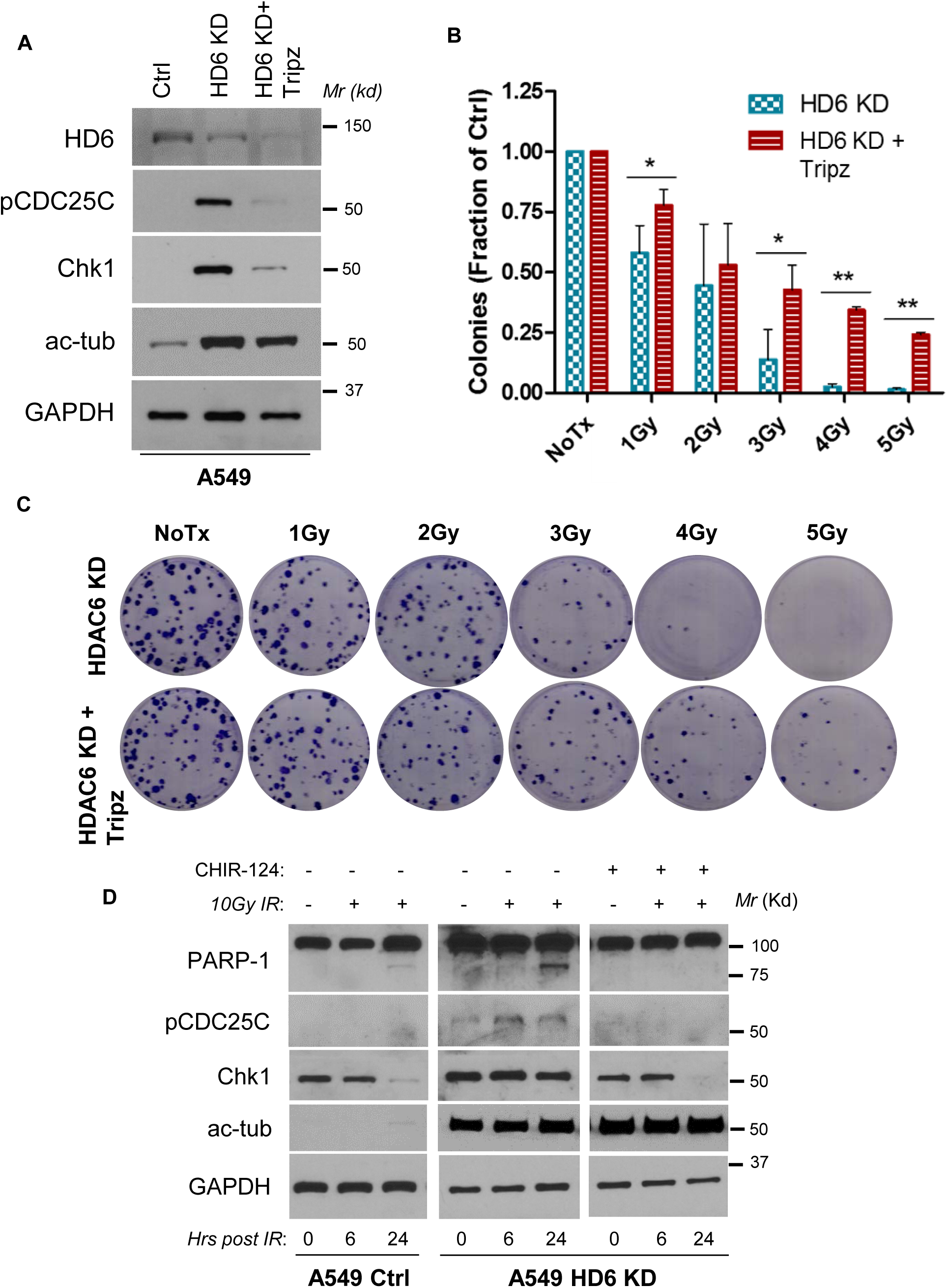
Radiosensitivity of HDAC6 knockdown cells is dependent on Chk1 activity. **A.)** A549 HDAC6 knockdown cells were transfected with a TRIPZ inducible lentiviral shRNA-expressing plasmid against Chk1, creating the Chk1Tripz line that is HDAC6 and Chk1 knocked-down. **B.)** Chk1Tripz and A549 HDAC6 knockdown cells were plated in triplicate at a concentration of 150 cells/well and treated with the indicated dose of radiation. Cells were incubated for 12 days, fixed with crystal violet, and quantified. Single sample t test, *p<0.05, **p<0.0008. **C.)** Representative images of A549 HDAC6 stable knockdown and Chk1Tripz colony formation assays described in (C). **D.)** A549 control and HDAC6 stable knockdown cells were pre-treated with 0.25μM of potent Chk1 inhibitor CHIR-124 prior to 10Gy irradiation. At the indicated timepoints, cells were harvested and probed for the indicated proteins via western blot.

To test the impact of pharmacological inhibition of Chk1 on the radiosensitivity of NSCLC cells, we utilized CHIR-124, a novel and potent Chk1 inhibitor with 2,000-fold less activity against Chk2 and 500-5,000-fold less activity against CDK2/4 and Cdc2 (37). We chose this inhibitor in lieu of clinically relevant inhibitor UCN-01 (38) due to the need for Chk1-specific inhibition over a kinetic profile suitable for use in humans. We pre-treated A549 control and HDAC6 knockdown cells with either CHIR-124 or DMSO (vehicle) for 24 hours prior to treatment with 10 Gy radiation, and found that the combination of Chk1 inhibition and radiation had an additive effect in control cells, with a faint PARP-1 cleavage band detectable 24 hours after radiation alone. In contrast, PARP-1 cleavage was detectable 24 hours after radiation alone in the HDAC6 knockdown cells, but pre-treatment with CHIR-124 protected these cells from radiation-induced PARP-1 cleavage [Figure 7D, right two panels]. Notably, total Chk1 protein levels in HDAC6 knockdown cells receiving combination treatment resolved in a manner similar to what is observed in control cells receiving either treatment scheme. This may suggest that the active variant of Chk1 is the most stable in HDAC6 deficient cells, as Chk1 inhibited at its catalytic domain loses the exquisite stability seen in cells deprived of total HDAC6 activity. Clarity concerning the full mechanism of HDAC6’s interaction with and control of Chk1 activity will require further study.

### Expression of active Chk1 in the nucleus is associated with NSCLC patient overall survival

To explore the indicative value of active Chk1 in NSCLC, we used a cohort of 187 early-stage NSCLC patients treated by surgery alone (28). Cohort demographic and clinical characteristics are summarized in Table 1. We determined the expression of pS317 Chk1 by Automated Quantitative Analysis (AQUA) as described in the Materials and Methods. As shown in Figure 8 and Tables 2 and 3, we have found that a higher level of pS317Chk1 is associated with a longer overall survival in this cohort with p=0.008. This clinical data supports our hypothesis that active Chk1 in the nucleus would execute proper checkpoint control and lead to favorable outcomes from patients. Further investigations examining another active Chk1 (pS345) and using a larger cohort including advanced NSCLC patients received ionizing radiation and chemotherapy are warranted.

**Table 2.** Result of multivariable cox model for OS.

| **Variable** | **n** | **HR (95% CI)** | **P value** |
| --- | --- | --- | --- |
| **p317-Chk1** |  |  |  |
| High | 102 | Ref. |  |
| Low | 67 | 1.66 (1.03-2.68) | **0.037** |
| **Sex** |  |  |  |
| Female | 80 | Ref. |  |
| Male | 89 | 1.70 (1.02-2.84) | **0.044** |
| **Age** | 169 | 1.03 (1.00-1.06) | **0.041** |
| **Stage** |  |  |  |
| IA | 76 | Ref. |  |
| IB | 93 | 1.45 (0.89-2.36) | 0.135 |
| **Histology** |  |  |  |
| Adeno or BAC | 88 | Ref. |  |
| Squamous | 62 | 0.60 (0.34-1.05) | 0.072 |
| L or LNE | 19 | 1.31 (0.66-2.60) | 0.435 |
| **Smoking** |  |  |  |
| Never | 11 | Ref. |  |
| Quit | 112 | 2.66 (0.62-11.49) | 0.190 |
| Active | 46 | 4.57 (1.03-20.22) | **0.045** |
| **PS** |  |  |  |
| 0 | 122 | Ref. |  |
| 1 | 47 | 2.04 (1.20-3.47) | **0.009** |

**Table 3.** Association between p317-Chk1 expression and clinicopathologic features.

| **Variable** | **High (n=109)** | **Low (n=77)** | **P value\*** |
| --- | --- | --- | --- |
| **Sex - no. (%)** |  |  | 0.459 |
| Female | 53 (49) | 33 (43) |  |
| Male | 56 (51) | 44 (57) |  |
| **Stage - no. (%)** |  |  | 0.052 |
| IA | 56 (51) | 28 (36) |  |
| IB | 53 (49) | 49 (64) |  |
| **Histology - no. (%)** |  |  | 0.403 |
| Adeno or BAC | 57 (52) | 39 (51) |  |
| Squamous | 42 (39) | 26 (34) |  |
| L or LNE | 10 (9) | 12 (16) |  |
| **Smoking status - no. (%)** |  |  | 0.812 |
| Never | 7 (6) | 4 (5) |  |
| Quit | 65 (60) | 48 (62) |  |
| Active | 30 (28) | 18 (23) |  |
| Missing | 7 (6) | 7 (9) |  |
| **PS - no. (%)** |  |  | 0.165 |
| 0 | 81 (74) | 47 (61) |  |
| 1 | 24 (22) | 23 (30) |  |
| Missing | 4 (4) | 7 (9) |  |
| **Age - median (range)** | 70.7 (47,84.1) | 70.2 (45.9,84.4) | 0.728 |

\*Fisher's exact test or Wilcoxon test as appropriate

**Figure 8:**
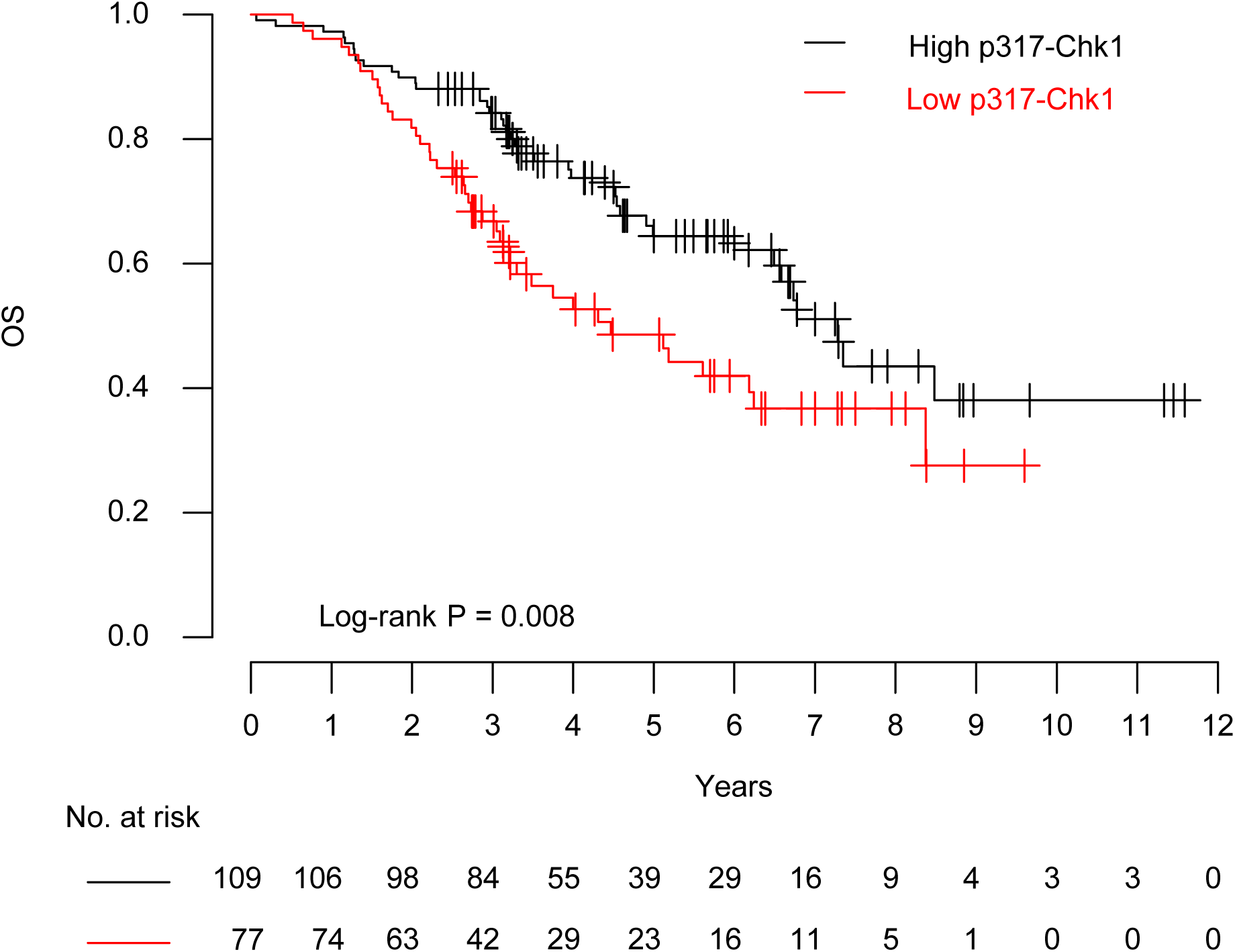
High levels of pChk1 S317 are associated with increased overall survival in NSCLC patients. Kaplan-Meier curve, univariate analysis of the overall survival of 187 NSLCL patients stratified by p S317 Chk1 status, with this status determined via AQUA staining.

## Discussion

Here, we describe a novel mechanism through which HDAC6 depletion sensitizes NSCLC to ionizing radiation-induced cell death. We have identified a mediator of this sensitization, Chk1, as a potential target of HDAC6’s E3 ubiquitin ligase activity. Chk1 is a critical Ser/Thr kinase central to the maintenance of genomic integrity, activated in response to single- and double-strand DNA breaks by ATR (39) and, to a lesser extent, ATM (40). Active Chk1 phosphorylates numerous downstream targets to arrest the cell cycle in S/G2/M, allowing time for DDR proteins to repair the initiating damage. Due to its central role in genomic preservation Chk1 is classically considered to be tumor-promoting, and clinical trials targeting Chk1 are ongoing. However, these trials haven’t proceeded past phase II due to multiple factors, including a lack of antitumor efficacy, excessive toxicity, and a concomitant activation of compensatory ATM and ERK1/2 signaling (41). Chk1 loss is embryonic lethal in mice (18), so even as increasingly specific inhibitors are developed Chk1 activity may prove too broad for this treatment direction to achieve FDA approval. In light of these failures in the clinic, multiple groups have suggested an alternative method of targeting Chk1; swinging the pendulum in the opposite direction and constitutively activating Chk1 (17).

Spearheaded by Dr. Tony Hunter and Dr. Youwei Zhang, this alternative Chk1 targeting strategy is the result of major insights made in Chk1 spatio-temporal regulation. In the nucleus, active Chk1 can arrest the cell cycle and activate homologous recombination proteins Rad51 and BRCA2. However, phosphorylation on S345 can also target Chk1 for nuclear export via Crm1 (42), inactivation by phosphatase PP2A (43), and ubiquitination by nuclear E3 ubiquitin ligase CTD2 (21). The fraction of Chk1 that is exported to the cytosol is still active, but in this compartment it can only perpetuate cell cycle arrest (44). Cytosolic Chk1 can also be ubiquitinated by E3 ubiquitin ligase Fbx6 (19), which allows for relief of arrest and the cell to enter mitosis. While this model of Chk1 regulation has been widely accepted, it does not necessarily conflict with our discovery of a new Chk1 E3 ligase. Tumor suppressor p53 has over 10 identified E3 ligases (23), and having multiple regulators fits the need for the cell to have tight and highly regulated control of DNA damage responders.

Despite these groups acknowledging that constitutive Chk1 activation may be detrimental to cancer cell viability, our group is the first to propose a method for accomplishing this aim. In multiple cell lines, we have demonstrated that HDAC6 knockdown increases basal Chk1 levels [Figure 3], and that further treatment of these cells with DNA damaging agents exacerbates Chk1 activation and persistence in parallel to their inability to resolve breaks [Figure 3]. The amount of damage HDAC6 knockdown cells accumulate, as indicated by γ-H2AX foci, is significant in both its intensity and duration when compared to the control counterparts [Figure 2]. Taken together, the data presented in this study supports the following mechanism of Chk1 regulation: upon recognition of radiation-induced damage, nuclear Chk1 is activated by upstream kinases and perpetuates cell cycle arrest and DNA repair. Active Chk1 is then exported to the cytosol, where it continues to arrest the cell cycle and allows time for the now active DNA repair factors to resolve the damage. Upon recognition of Chk1, we hypothesize HDAC6 will ubiquitinate and target Chk1 for degradation, releasing the cell from arrest and allowing mitosis to occur. This model, and in particular the potential clinical impact of elevated active Chk1 on tumors, is bolstered by our analysis of NSCLC patients stratified by Chk1 activation [Figure 8]. We observe that patients with higher active Chk1 in the nucleus (pChk1 on Serine 317) had significantly higher long-term survival than patients that exhibited lower levels of active Chk1, in a cohort of surgically resected patients with stage I NSCLC. HDAC6 protein levels are elevated in NSCLC tumors when compared to control tissues, so this model may explain one mechanism behind the rapid growth of NSCLC. Conversely, inhibition of HDAC6’s E3 ubiquitin ligase activity could eliminate a mechanism NSCLC is reliant upon, inducing a perpetual cell cycle arrest in response to ionizing radiation and forcing the cell into mitotic catastrophe and subsequent cell death. Further research is necessary to assess the voracity of this mechanism.

While E3 ligase inhibitors are commercially available as research tools, the majority are nonspecific due to the ubiquity of the pathways E3 ligases participate in, and thus there is virtually no interest in translating this strategy into the clinic. Unfortunately, we cannot circumvent this shortcoming with clinically relevant HDAC6-specific inhibitors, as both ACY-1215 and ACY-241 are designed to solely inhibit the deacetylase activity of the DAC2 domain. Our research supports the design of an inhibitor able to target the DAC1 domain of HDAC6, or potentially both the DAC1 and DAC2 domain, to achieve total HDAC6 inhibition in vivo. Development of such an inhibitor, if truly HDAC6-specific and fitting the patient tolerability seen with HDAC6 deacetylase inhibitors, would provide a strategy to sensitize non-small cell lung cancer patients to conventional DNA damaging agents. Alternatively, methods to target the upstream regulators of HDAC6 could indirectly decrease its protein levels and reduce deacetylase and E3 ubiquitin ligase activity. For instance, tamoxifen treatment of MCF-7 breast cancer cells prevented estradiol-stimulated HDAC6 accumulation and α-tubulin deacetylation (45), and nitric oxide (NO) is able to increase the acetylation of α-tubulin in in A549 cells by preventing the essential S-nitrosylation of HDAC6 (46). More directly, Cullin 3SPOP has been reported to destabilize HDAC6 via polyubiquitination, and this interaction has suggested that SPOP serves a tumor suppressor function through this mechanism (47). Further research is needed to identify the full spectrum of proteins that regulate HDAC6 stability, especially due to the relatively long half-life of HDAC6 in vivo, but the advantage of this approach is that strategies to modulate HDAC6 enzymatic activity already exist and can be tested to determine whether the DAC1 E3 ubiquitin ligase activity is being inhibited alongside the DAC2 deacetylase activity.

## Acknowledgements

We would like to thank the Wayne State MICR core for assistance with cell sorting, flow cytometry, and subsequent data analysis. We are grateful for the support from Dr. Youwei Zhang, who provided insights of this project as well as a series of Chk1 deletion mutants, and a careful reading of this manuscript. We would also like to acknowledge Dr. Patrick Matthias, the creator of the HDAC6 KO transgenic mouse model, and Dr. Eduardo M. Sotomayor, who transferred HDAC6 KO transgenic mice with permission from Dr. Matthias.

